# ABCB7 deficiency increases endoplasmic reticulum–mitochondria coupling and mTOR-independent autophagy flux in H9c2 cardiomyocytes

**DOI:** 10.1101/2023.08.18.553927

**Authors:** Shivanshu Chandan, Ganesh Kosher

**Affiliations:** Department of Pharmaceutical Sciences, Central University of Haryana, Mahendergarh, Haryana, India 123031

**Keywords:** ABCB7, Mitochondria, ER, Cardiac hypertrophy, Heart failure

## Abstract

ABCB7 deficiency during chronic cardiac hypertrophy contribute to mitochondrial dysfunction, metabolic shift and worsens cardiac function. Here, we explored that ABCB7 deficiency contribute to tethering of mito-ER and in turn mitochondrial dysfunction in H9c2 cells. We also investigated the mechanistic link between mitochondrial dysfunction and ABCB7 deficiency in these cells. Knockdown of ABCB7 was performed by siABCB7 plasmids or control vectors using lipofectamine 2000. To rescue the changes produced by siABCB7, ABCB7 overexpression was performed using ABCB7 overexpression vector. After knockdown or overexpression, cells were harvested for transmission electron microscopy (TEM), RT-PCR or Immunofluorescence analysis. Knockdown of ABCB7 in H9C2 cells resulted in enhanced tethering of mito.-ER contact sites, and increased mito.-ER distance. To our surprise, the downregulation of ABCB7 did not alter the cristae structure or morphology in these cells. On the mechanistic front, Knockdown of ABCB7 in H9C2 cells MTOR-independent AMPK-dependent macroautophagic/autophagic flux. ABCB7 downregulation did not result in cell death in these cells; this phenomenon could work independent of cell death in H9c2 cells.

## Introduction

The ATP-binding cassette (ABC) sub-family of protein tightly regulate mitochondrial function^1-5^. The ABCB7 gene encodes a protein known as the ATP-binding cassette sub-family B member 7, which is also commonly referred to as ABCB7. This protein is a crucial component of cellular processes related to iron metabolism and homeostasis^8-12^. ABCB7 functions as a transporter located in the inner mitochondrial membrane, where it plays a significant role in the transport of iron-sulfur clusters, essential components of various proteins involved in electron transfer and metabolic reactions^13-16^.

The primary function of ABCB7 is to facilitate the transport of iron-sulfur clusters from the mitochondria to the cytoplasm, where they are utilized for the assembly of various iron-sulfur proteins. These proteins are essential for several cellular functions, including DNA synthesis, energy production, and various enzymatic reactions^12-16^. Additionally, ABCB7 also plays a role in exporting excess iron from the mitochondria, preventing the accumulation of toxic levels of iron within the organelle^5-10^.

Mutations or dysregulation in the ABCB7 gene can lead to various disorders collectively known as X-linked sideroblastic anemias (XLSA). These conditions are characterized by a disruption in the production of healthy red blood cells due to impaired iron metabolism. Individuals with XLSA may exhibit symptoms such as anemia, fatigue, and other complications stemming from insufficient red blood cell production^15-17^.

In the context of the heart, ABCB7 (ATP-binding cassette sub-family B member 7) serves a vital role in maintaining cellular health and function through its involvement in iron metabolism. Iron is a crucial element for various processes in the heart, including energy production, oxygen transport, and overall cardiac function^15-21^. ABCB7 is primarily found in the mitochondria of cardiac cells, where it functions as a transporter responsible for the movement of iron-sulfur clusters. These clusters are essential components of proteins involved in electron transport chains and enzymatic reactions critical for generating energy in the form of adenosine triphosphate (ATP)^22-25^.

The heart has high energy demands due to its continuous pumping action, and a well-functioning iron metabolism is crucial for meeting these energy needs. ABCB7 plays a key role in ensuring that iron-sulfur clusters are correctly transported within the mitochondria, where they are incorporated into important proteins involved in cellular respiration and oxidative phosphorylation. These processes are fundamental for producing ATP, the energy currency of cells^15, 17, 25^.

Disruptions in ABCB7 function can lead to imbalances in iron homeostasis within cardiac cells. Iron overload or deficiency can adversely affect cardiac function, leading to conditions such as cardiomyopathies, which are characterized by structural and functional abnormalities in the heart muscle. Proper ABCB7 function is crucial for preventing iron-related oxidative stress and maintaining the overall health of cardiac cells^17-25^.

In summary, ABCB7’s role in the heart involves facilitating the transport of iron-sulfur clusters within mitochondria, thereby contributing to energy production and overall cardiac function. Maintaining a delicate balance in iron metabolism through ABCB7 is vital for preventing cardiac complications and ensuring the heart’s efficient performance. Here, we explored that ABCB7 deficiency contribute to tethering of mito-ER and in turn mitochondrial dysfunction in H9c2 cells. We also investigated the mechanistic link between mitochondrial dysfunction and ABCB7 deficiency in these cells.

## Results

In an earlier study, Vikas Kumar et al., identified ABCB7’s role in the heart and discovered that deficiency of this gene in the heart can lead to mitochondrial dysfunction. Here, we knockdown the ABCB7 gene in H9C2 cells using siABCB7 plasmid and harvested the cells for transmission electron microscopy (TEM). We analyzed the cells for mitochondria and ER structure, morphology, tethering and contact site analysis using software ImageJ. Mito-ER contract sites were increased in the ABCB7 silenced H9c2 cells. The TEM images were quantified and revealed that ER distance was decreased in ABCB7 silenced h9c2 cells. The mitochondrial structure, and ER structure were however unaltered by silencing of ABCB7 in these cells. These results suggest a tethering of these two organelles on ABCB7 silencing without affecting their morphology. We in addition, also observed that the cristae no., its morphology and size were not changed on ABCB7 silencing (Fig.1A-Fig.1D).

**Figure 1.**
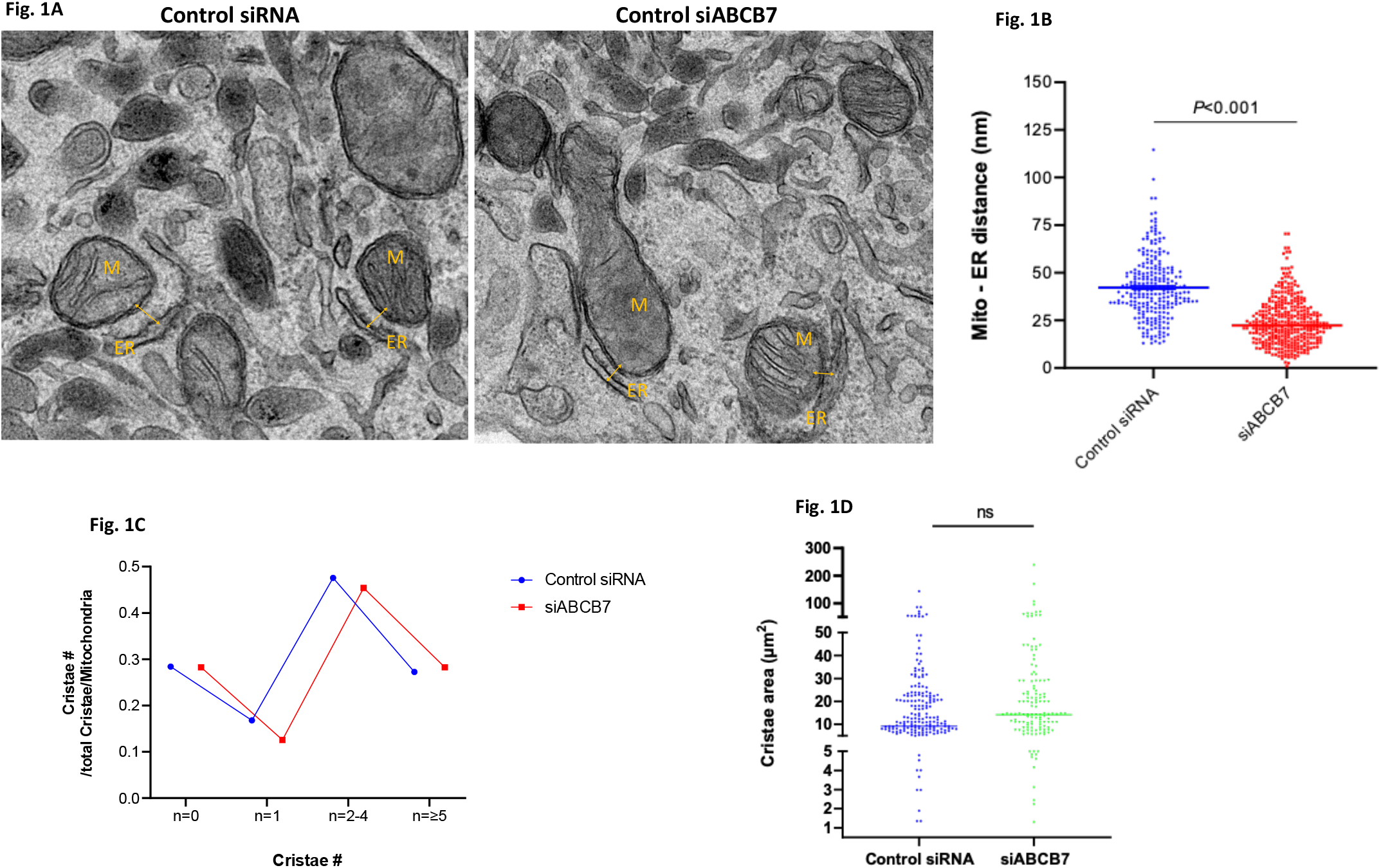

The increased contact sites between Mito-ER allowed the disturbed and decreased calcium ion influx across the mitochondrial membrane into the mitochondria. The calcium ion flux rate and peak calcium concentration were lower in the ABCB7 silenced H9c2 cells (Fig. 2A, 2B).

**Figure 2.**
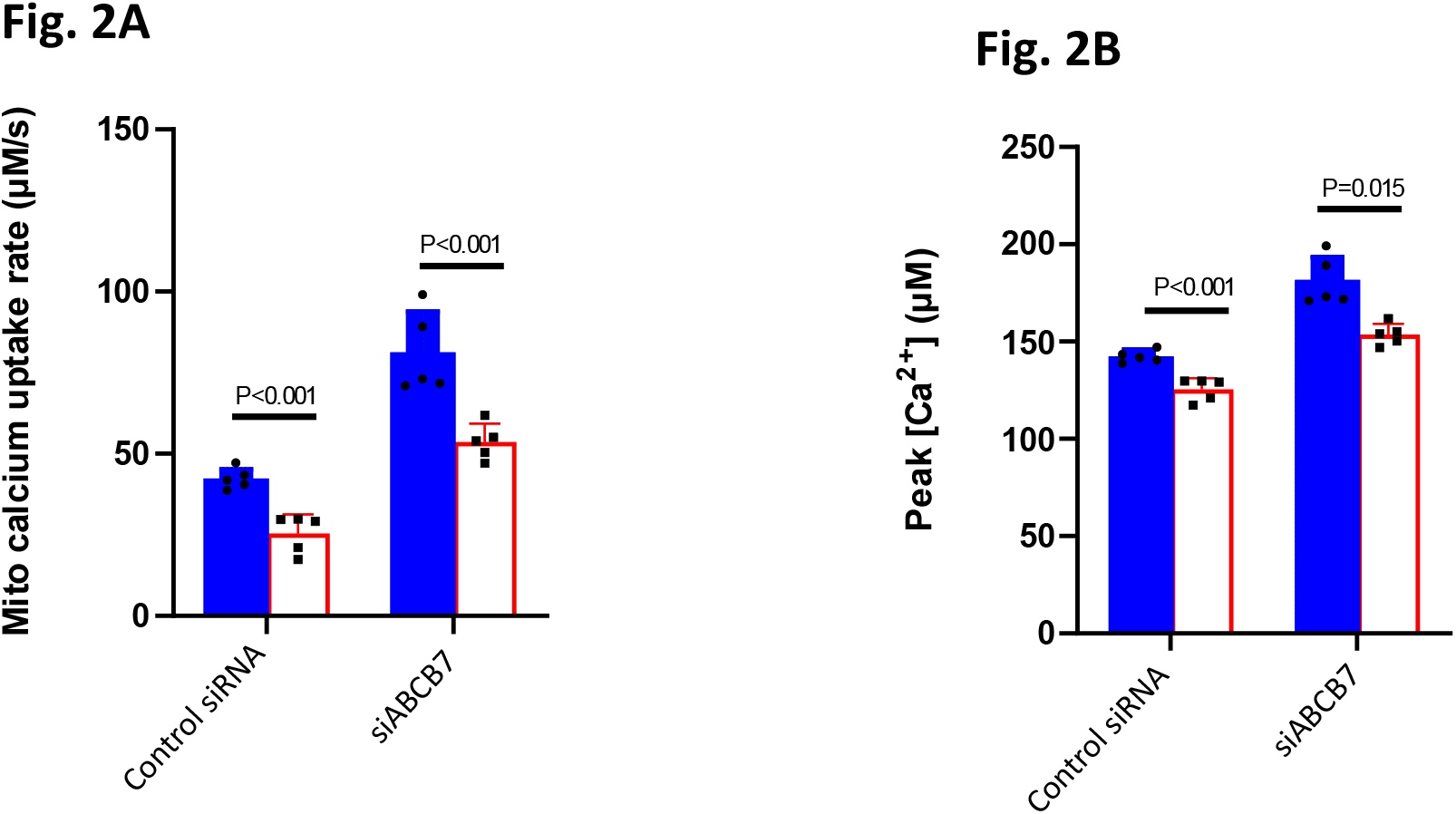

On the mechanistic front, ABCB7 silencing increased AMPK mediated autophagy flux across the mitochondrial membrane (data not shown).

In conclusion, these results suggest that ABCB7 is crucial for mitochondrial homeostasis and in turn metabolic flux in the heart. It is still unknown if the ABCB7 deficiency contribute to inflammatory signals in the heart

## Discussion

ABCB7 (ATP-binding cassette sub-family B member 7) plays a significant role in cardiomyocytes, which are the specialized muscle cells that make up the heart. Its primary function in these cells is closely tied to iron metabolism, which is essential for various processes critical to cardiac function.

Iron-Sulfur Cluster Biogenesis: One of the most important roles of ABCB7 in cardiomyocytes is its involvement in the biogenesis of iron-sulfur clusters. Iron-sulfur clusters are co-factors essential for the function of numerous proteins involved in electron transport chains, enzyme activities, and redox reactions. These processes are crucial for energy production, particularly in the form of ATP through oxidative phosphorylation. ABCB7 helps transport the necessary iron and sulfur components for the assembly of these clusters within mitochondria, where they are integrated into proteins critical for energy generation.

Mitochondrial Function: Cardiomyocytes have high energy demands due to their continuous contractile activity. Proper mitochondrial function is essential to meet these energy requirements. Iron-sulfur proteins are key components of the electron transport chain, the process by which mitochondria generate ATP. ABCB7’s role in transporting iron-sulfur clusters ensures the correct assembly of these proteins, promoting efficient oxidative phosphorylation and ATP production.

Protection Against Oxidative Stress: Iron, while essential, can also be potentially harmful when present in excess. ABCB7 helps maintain iron homeostasis within cardiomyocytes, preventing iron overload and subsequent oxidative stress. Oxidative stress occurs when there is an imbalance between reactive oxygen species (ROS) production and the cell’s antioxidant defenses. By facilitating proper iron metabolism, ABCB7 contributes to reducing the risk of oxidative damage to cardiomyocytes, which is crucial for maintaining cardiac health.

Role in Cardiomyopathies: Dysregulation of iron metabolism in cardiomyocytes can lead to cardiomyopathies, which are conditions characterized by structural and functional abnormalities in the heart muscle. Mutations or malfunction of ABCB7 can disrupt iron-sulfur cluster biogenesis, leading to impaired mitochondrial function and energy production. This, in turn, can contribute to the development of cardiomyopathies and heart failure.

In summary, ABCB7’s role in cardiomyocytes is centered on facilitating proper iron-sulfur cluster biogenesis, mitochondrial function, and iron homeostasis. These functions are essential for meeting the high energy demands of the heart and preventing oxidative stress. Dysfunction of ABCB7 can have significant implications for cardiac health and contribute to the development of cardiac disorders.

In conclusion, these results suggest that ABCB7 is crucial for mitochondrial homeostasis and in turn metabolic flux in the heart. It is still unknown if the ABCB7 deficiency contribute to inflammatory signals in the heart

## Acknowledgment

We thank the Central University of Haryana for providing the facilities for this study.

## References

1. Hogue, D. L., Liu, L. & Ling, V. Identification and characterization of a mammalian mitochondrial ATP-binding cassette membrane protein. J. Mol. Biol. 285, 379–89 (1999).

2. Zutz, A., Gompf, S., Schägger, H. & Tampé, R. Mitochondrial ABC proteins in health and disease. Biochim Biophys Acta - Bioenerg. 1787, 681–90 (2009).

3. Richardson, D. R. et al. Mitochondrial iron trafficking and the integration of iron metabolism between the mitochondrion and cytosol. Proc Natl Acad Sci. 107, 10775–10782 (2010).

4. Schumacher, T. & Benndorf, R. A. ABC Transport Proteins in Cardiovascular Disease— A Brief Summary. Molecules 22, E589 (2017).

5. Rakvács, Z. et al. The human ABCB6 protein is the functional homologue of HMT-1 proteins mediating cadmium detoxification. Cell. Mol. Life Sci. 76, 1–14 (2019).

6. Pondarré, C. et al. The mitochondrial ATP-binding cassette transporter Abcb7 is essential in mice and participates in cytosolic ironsulfur cluster biogenesis. Hum. Mol. Genet. 15, 953–64 (2006).

7. Pondarre, C. et al. Abcb7, the gene responsible for X-linked sideroblastic anemia with ataxia, is essential for hematopoiesis. Blood 109, 3567–9 (2007).

8. Cavadini, P. et al. RNA silencing of the mitochondrial ABCB7 transporter in HeLa cells causes an iron-deficient phenotype with mitochondrial iron overload. Blood 109, 3552–3559 (2007).

9. Miyake, A. et al. Mutation in the abcb7 gene causes abnormal iron and fatty acid metabolism in developing medaka fish. Dev. Growth differ. 50, 703–16 (2008).

10. Nikpour, M. et al. The transporter ABCB7 is a mediator of the phenotype of acquired refractory anemia with ring sideroblasts. Leukemia 27, 889–896 (2013).

11. Lerebours, A., To, V. V. & Bourdineaud, J. P. Danio rerio ABC transporter genes abcb3 and abcb7 play a protecting role against metal contamination. J. Appl. Toxicol. 36, 1551–1557 (2016).

12. Li, J. & Cowan, J. A. Glutathione-coordinated [2Fe–2S] cluster: a viable physiological substrate for mitochondrial ABCB7 transport. Chem. Commun. 51, 2253–5 (2015).

13. Protasova, M. S. Whole-genome sequencing identifies a novel ABCB7 gene mutation for X-linked congenital cerebellar ataxia in a large family of Mongolian ancestry. Eur. J. Hum. Genet. 24, 550–555 (2016).

14. Dolatshad, H. Cryptic splicing events in the iron transporter ABCB7 and other key target genes in SF3B1-mutant myelodysplastic syndromes. Leukemia 30, 2322–2331 (2016).

15. Kumar, V., Sanawar, R., Jaleel, A., Santhosh Kumar, T.R. and Kartha, C.C., 2019. Chronic pressure overload results in deficiency of mitochondrial membrane transporter ABCB7 which contributes to iron overload, mitochondrial dysfunction, metabolic shift and worsens cardiac function. Scientific reports, 9(1), p. 13170.

16. Kumar, V., Santhosh Kumar, T.R. and Kartha, C.C., 2019. Mitochondrial membrane transporters and metabolic switch in heart failure. Heart failure reviews, 24, pp. 255–267.

17. Tarasov, K.V., Chakir, K., Riordon, D.R., Lyashkov, A.E., Ahmet, I., Perino, M.G., Silvester, A.J., Zhang, J., Wang, M., Lukyanenko, Y.O. and Qu, J.H., Vikas Kumar., 2022. A remarkable adaptive paradigm of heart performance and protection emerges in response to marked cardiac-specific overexpression of ADCY8. Elife, 11, p. e80949.

18. Kumar, V., Kumar, A.A., Joseph, V. et al. Untargeted metabolomics reveals alterations in metabolites of lipid metabolism and immune pathways in the serum of rats after long-term oral administration of Amalaki rasayana. Mol Cell Biochem 463, 147–160 (2020).

19. Mohan, N., Kumar, V., Kandala, D.T., Kartha, C.C. and Laishram, R.S., 2018. A splicing-independent function of RBM10 controls specific 3′ UTR processing to regulate cardiac hypertrophy. Cell reports, 24(13), pp. 3539–3553.

20. Rai, A., Kumar, V., Jerath, G., Kartha, C.C. and Ramakrishnan, V., 2021. Mapping drug-target interactions and synergy in multi-molecular therapeutics for pressure-overload cardiac hypertrophy. NPJ systems biology and applications, 7(1), p. 11.

21. Nair, R.S., Soghan, P.K., Shenoy, S.J., Prabhu, M.A., Kumar, V., Ramachandran, S. and Anilkumar, T.V., 2023. Mitigation of Fibrosis after Myocardial Infarction in Rats by Using a Porcine Cholecyst Extracellular Matrix. Comparative Medicine.

22. Pradhan, T., Kumar, V., Surya H E., Krishna, R., John, S., Jissa, V.T., Anjana, S., Chandramohan, K. and Nair, S.A., 2021. STIL endows oncogenic and stem-like attributes to colorectal cancer plausibly by Shh and Wnt signaling. Frontiers in Oncology, 11, p. 581671.

23. Kumar, V., Bermea, K.C., Kumar, D., Singh, A., Verma, A., Kaileh, M., Sen, R., Lakatta, E.G. and Adamo, L., 2023. Cardiomyocyte-specific adenylyl cyclase type-8 overexpression induces cell-autonomous activation of RelA and non-cell-autonomous myocardial and systemic inflammation. bioRxiv, pp. 2023–07.

24. Kumar, V., Aneesh, K.A., Kshemada, K., Ajith, K.G., Binil, R.S., Deora, N., Sanjay, G., Jaleel, A., Muraleedharan, T.S., Anandan, E.M. and Mony, R.S., 2017. Amalaki rasayana, a traditional Indian drug enhances cardiac mitochondrial and contractile functions and improves cardiac function in rats with hypertrophy. Scientific reports, 7(1), p. 8588.

25. Kwatra, M., Kumar, V., Jangra, A., Mishra, M., Ahmed, S., Ghosh, P., Vohora, D. and Khanam, R., 2016. Ameliorative effect of naringin against doxorubicin-induced acute cardiac toxicity in rats. Pharmaceutical biology, 54(4), pp. 637–647.

